# Impulsivity and risk-seeking as Bayesian inference under dopaminergic control

**DOI:** 10.1101/2020.10.06.327775

**Authors:** John G. Mikhael, Samuel J. Gershman

## Abstract

Bayesian models successfully account for several of dopamine (DA)’s effects on contextual calibration in interval timing and reward estimation. In these models, tonic levels of DA control the precision of stimulus encoding, which is weighed against contextual information when making decisions. When DA levels are high, the animal relies more heavily on the (highly precise) stimulus encoding, whereas when DA levels are low, the context affects decisions more strongly. Here, we extend this idea to intertemporal choice and probability discounting tasks. In intertemporal choice tasks, agents must choose between a small reward delivered soon and a large reward delivered later, whereas in probability discounting tasks, agents must choose between a small reward that is always delivered and a large reward that may be omitted with some probability. Beginning with the principle that animals will seek to maximize their reward rates, we show that the Bayesian model predicts a number of curious empirical findings in both tasks. First, the model predicts that higher DA levels should normally promote selection of the larger/later option, which is often taken to imply that DA decreases ‘impulsivity,’ and promote selection of the large/risky option, often taken to imply that DA increases ‘risk-seeking.’ However, if the temporal precision is sufficiently decreased, higher DA levels should have the opposite effect—promoting selection of the smaller/sooner option (higher impulsivity) and the small/safe option (lower risk-seeking). Second, high enough levels of DA can result in preference reversals. Third, selectively decreasing the temporal precision, without manipulating DA, should promote selection of the larger/later and large/risky options. Fourth, when a different post-reward delay is associated with each option, animals will not learn the option-delay contingencies, but this learning can be salvaged when the post-reward delays are made more salient. Finally, the Bayesian model predicts correlations among behavioral phenotypes: Animals that are better timers will also appear less impulsive.

## Introduction

The neuromodulator dopamine (DA) has been repeatedly associated with choice impulsivity, the tendency to prioritize short-term over long-term reward. Impulsive behaviors characterize a number of DA-related psychiatric conditions [1], such as attention-deficit/hyperactivity disorder [2–6], schizophrenia [7, 8], addiction [9, 10], and dopamine dysregulation syndrome [11, 12]. Furthermore, direct pharmacological manipulation of tonic DA levels in humans [13, 14] and rodents [15, 16] has corroborated a relationship between DA and impulsivity. The standard approach to measuring impulsive choice is the intertemporal choice task (ITC), in which subjects choose between a small reward delivered soon and a large reward delivered later [17]. A subject’s preference for the smaller/sooner option is often taken as a measure of their impulsivity, or the extent to which they discount future rewards [18–21].

In the majority of animal studies, higher DA levels have been found to promote selection of the larger/later option (inhibiting impulsivity) [15, 22–28]. However, the inference that DA agonists inhibit impulsivity has been challenged in recent years, in part because, when ITCs are administered to humans, DA agonists seem to *promote* impulsivity [29]. Perhaps relevant to this contrast is that, while impulsive choices in humans are assessed through hypothetical situations (‘Would you prefer $1 now or $10 in one month?’), ITCs in animals more closely resemble reinforcement learning tasks involving many trials of experienced rewards and delays. Complicating this picture further, the effect of DA, even within animal studies, is not straightforward. While in most studies, DA appears to decrease impulsivity, DA has been found to systematically increase impulsivity under some conditions [30–32], such as when the delay period is uncued [16] or when different delays for the larger/later option are presented in decreasing order across training blocks [33].

The relationship of DA with impulsive choice finds a parallel in its relationship with risk-seeking. Disruptions in risk preferences feature prominently in a number of DA-related conditions [1], including Parkinson’s disease [34–37], schizophrenia [38, 39], and attention-deficit/hyperactivity disorder [40]. Moreover, direct manipulation of DA levels in Parkinson’s patients [41], healthy humans [42], and rodents [29] has further established a link between DA and risk preferences.

Risk-seeking can be formalized as the tendency to prioritize uncertain rewards over less uncertain rewards of equal average value. For example, a risk-seeker will preferentially select an option yielding a reward of magnitude 10 on 50% of trials and no reward in the remaining trials, over an option yielding a reward of magnitude 5 on 100% of trials. A standard measure of risk-seeking is the probability discounting task (PD) [43–45], where subjects choose between a small reward delivered with complete certainty and a large reward that is only delivered with some probability. Subjects that are more likely to select the large/risky option than other subjects—regardless of reward probability—are labeled as being more risk-seeking. Though studies involving direct pharmacological manipulation have highlighted a key role for DA in setting this preference, the directionality of DA’s effect has remained unclear: Whereas St Onge and Floresco [46] have found that increasing the DA level promotes risk-seeking in PDs, follow-up work has shown that DA may have exactly the opposite effect, depending on how the training blocks are ordered [47], a variable whose relevance for risk preferences is not immediately obvious.

Animal behavior in ITCs and PDs can be reinterpreted from a reinforcement learning perspective. With repeated trials of the same task, an optimal agent can learn to maximize its total accumulated rewards by estimating the reward rate for each option (reward magnitude divided by total trial duration) and choosing the option with the higher reward rate. Thus if the larger/later option has a sufficiently large reward or sufficiently short delay, it will be the optimal choice. However, if its reward were sufficiently small or its delay sufficiently long, the smaller/sooner option may be the superior choice instead, without any assumption of ‘discounting.’ Under this view, animals do not necessarily discount rewards at all, but rather make choices based on a reward-rate computation. The notions of true impulsivity in ITCs and risk-seeking in PDs have persisted, however, because animals tend to choose the smaller/sooner and large/risky options even when they objectively yield fewer rewards over many trials.

To address the question of whether animals compare reward rates, a body of theoretical and experimental work focused on impulsivity has demonstrated that the suboptimal tendency to choose the smaller/sooner option is better explained by *temporal* biases than by biases of choice [48–50] (see also [51]). This work has shown that animals behave in a way consistent with maximizing their reward rates, but they underestimate the elapsed time—and in particular, the periods after receiving the reward and before beginning the next trial. Thus animals estimate the reward rates for each option based largely on the pre-reward delays. This bias disproportionately benefits the smaller/sooner option, which has a much shorter pre-reward delay. As a result, the animals make choices that can be interpreted as impulsive. Said differently, animals disproportionately underestimate the total trial duration for the smaller/sooner option compared to the larger/later option, making the former more appealing. While this discounting-free view derives animal behavior from a normative framework (maximizing reward rates), how and why DA modulates choice preferences remains the subject of much speculation.

In this paper, we build on recent theoretical work that cast DA in a Bayesian light [52, 53]. Here, DA controls the precision with which cues are internally represented, which in turn controls the extent to which the animal’s estimates of the cues are influenced by context. In Bayesian terms, which we discuss below, DA controls the precision of the likelihood relative to that of the prior (the context). This framework predicts a well-replicated result in the interval timing literature, referred to as the ‘central tendency’ effect: When temporal intervals of different lengths are reproduced under DA depletion (e.g., in unmedicated Parkinson’s patients), shorter intervals tend to be overproduced and longer intervals tend to be underproduced, and DA repletion rescues accurate timing [54–56]. We recently extended this framework to the representation of reward estimates [57]. In this case, the Bayesian framework predicts that DA should tip the exploration-exploitation balance toward exploitation, in line with empirical findings [58–60] (but see [61, 62]).

We show here that, under the Bayesian theory, higher DA levels should promote behaviors consistent with lower impulsivity in the standard ITC (selection of the larger/later option), but should have the opposite effect when the temporal precision of the delay period is selectively and sufficiently reduced. In both cases, high enough levels of DA should elicit preference reversals, and not only an amplification of the current preference. Furthermore, in manipulations of temporal precision, if animals are more likely to select the larger/later option at baseline, DA administration will tend to reverse that preference (promote the smaller/sooner option), and vice versa. We show that animals should not learn the contingencies between options and their post-reward delays, but that this learning can be salvaged if the post-reward delays are made more salient. We show that animals that display more precise behaviors in interval timing tasks should also appear less impulsive. Finally, we reproduce this analysis for the case of risk-seeking and PDs: Depending on the relative balance between the uncertainty about the reward magnitude and the uncertainty about the reward probability, we show that DA can either promote or suppress selection of the large/risky option.

## Methods

### The Bayesian theory of dopamine

An agent wishing to encode information about some cue must contend with noise at every level, including the information source (which is seldom deterministic), storage (synapses are noisy), and signaling (neurons are noisy) [63]. We can formalize the noisy encoding as a mapping from an input signal (e.g., experienced reward) to a distribution over output signals (e.g., firing rates). For the purposes of this paper, we will remain agnostic about the specific neural implementation of the mapping, and instead discuss it in abstract terms. Thus a noisy encoding of some variable can be represented by a distribution over values: Tight distributions correspond to encodings with low noise (Fig. 1A), whereas wide distributions correspond to encodings with high noise (Fig. 1B).

**Figure 1:**
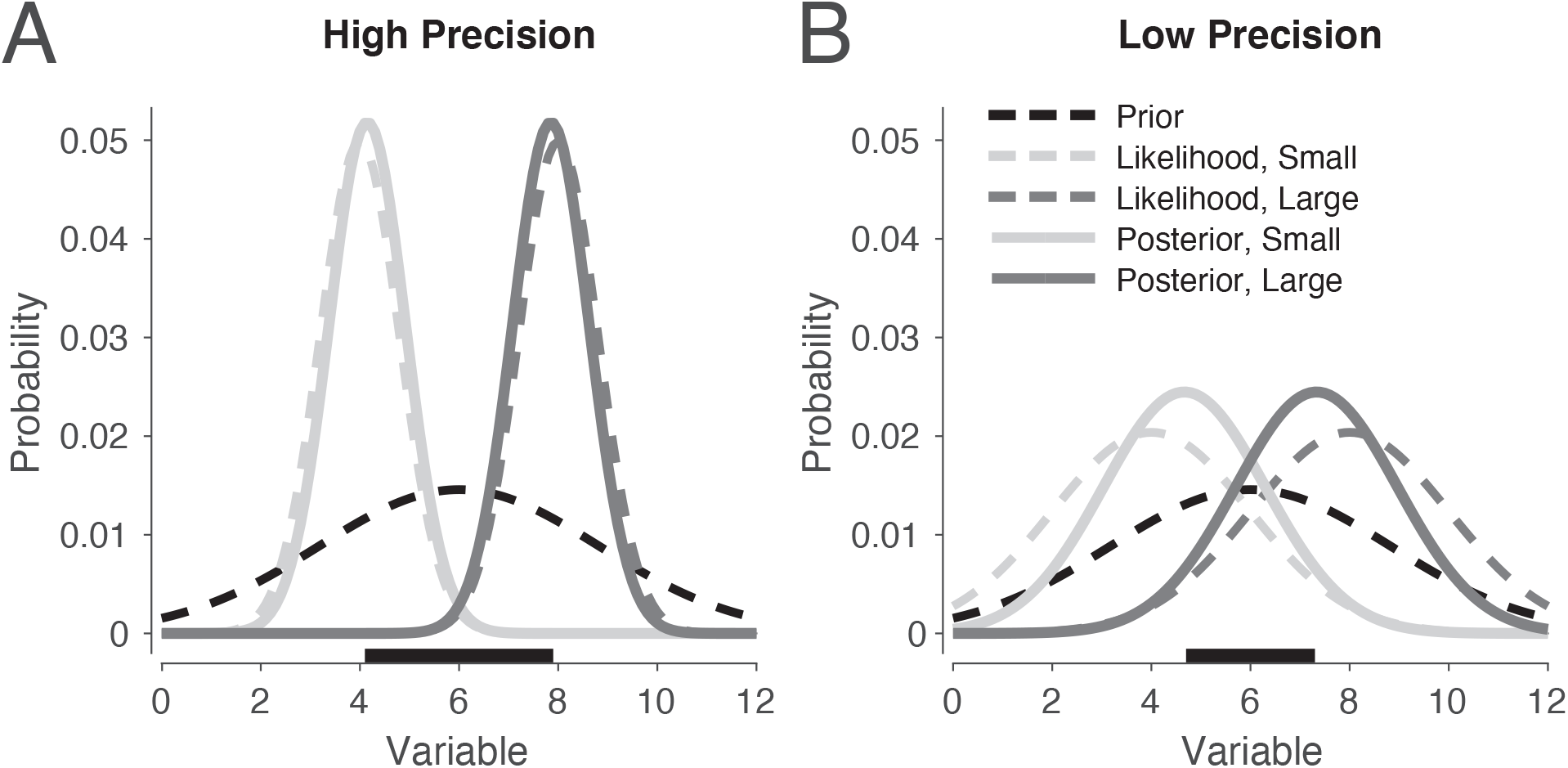
Contextual influence is stronger when the encoding precision is low. Distributions for two signals, one small and the other large. (A) When the encoding precision is high compared to the prior precision, the posteriors do not deviate significantly from the likelihood. (B) As the encoding precision decreases, the posteriors migrate toward the prior. The horizontal black segments illustrate the difference in posterior means under high vs. low precision.

Consider, then, a scenario in which an animal must estimate the average yield of a reward source from noisy samples. Because of the animal’s uncertainty about the average yield (the encoding distribution has non-zero spread), its final estimate can be improved by utilizing other sources of information. For example, if the nearby reward sources tend to yield large rewards, then the animal should form an optimistic estimate of the reward source’s average yield. Similarly, if nearby reward sources yield small rewards, then the animal should form a pessimistic estimate. Formally, the contextual information can be used to construct a prior distribution over average yield, and the encoding distribution can be used to construct a likelihood function for evaluating the consistency between the encoded information and a hypothetical average yield. Bayes’rule stipulates that the animal’s final probabilistic estimate should reflect the product of the likelihood and prior:

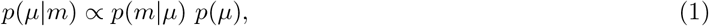

referred to as the posterior distribution. Here, *μ* is the variable being estimated (the reward yield), *m* is the stored value, *p*(*m*|*μ*) is the likelihood, and *p*(*μ*) is the prior. For simplicity, we take these distributions to be Gaussian throughout. Under standard assumptions for Gaussian distributions, the estimate 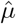 corresponds to the posterior mean:

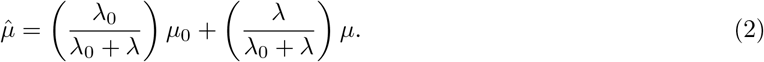

Here, *μ*_0_, *λ*_0_, *μ*, and *λ* represent the prior mean, prior precision, likelihood mean, and encoding precision, respectively. In words, the agent takes a weighted average of the prior mean *μ*_0_ and the likelihood mean *μ*— weighted by their respective precisions *λ*_0_ and *λ* after normalization—to produce its estimate, the posterior mean 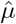. Intuitively, the tighter each distribution, the more it pulls the posterior mean in its direction.

The Bayesian theory of DA asserts that increasing the DA level increases the encoding precision *λ*, where the prior here represents the distribution of stimuli (i.e., the context). Thus when DA is high, the estimate 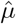 does not heavily depend on contextual information, whereas when it is low, Bayesian migration of the estimate to the prior is strong (compare Fig. 1A and B). Shi et al. [56] have applied this theory to interval timing and shown that it predicts DA’s effects on the central tendency: Parkinson’s patients who are on their medication will have high *λ*, qualitatively corresponding to Fig. 1A. Then the temporal estimates for the short and long durations will be very close to their true values (here, 4 and 8 seconds). On the other hand, patients who are off their medication will have low *λ*, corresponding to Fig. 1B. Thus the estimates for both durations will migrate toward the prior mean, or the average of the two durations. In other words, the estimate for the short duration will be overproduced, and the estimate for the long duration will be underproduced, as observed [54, 55].

The Bayesian model can also be applied to reward magnitudes [64, 65]. Imagine a bandit task in which an agent samples from two reward sources, one yielding small rewards on average and the other yielding large rewards on average. Under lower levels of DA, the central tendency should decrease the difference between the two reward estimates (compare lengths of black segments on the x-axis in Fig. 1A and B). Under standard models of action selection, animals are more likely to choose the large option when the difference between the two estimates is large, and become more and more likely to sample other options as the difference decreases (see next section). This means that lower levels of DA should promote selection of the smaller reward, often taken to indicate a drive to ‘explore,’ as empirically observed [58–60] (but see [61, 62]). Thus, as previously proposed, DA may be interpreted as controlling the exploration-exploitation trade-off [66]. This is in line with the ‘gain control’ theory of DA, in which high DA levels have been hypothesized to amplify the difference between reward estimates during decision making [66–70]. The Bayesian theory of DA subsumes the gain control view (see next section), but importantly, under this theory, animals do not become intrinsically more explorative or exploitative under different DA levels, but rather modify their behaviors to match the estimated difference in rewards.

We can also compare the degree of the central tendency in temporal and reward estimation, which will be important in the Results. Empirically, the central tendency in temporal tasks is normally weak. While it can be unmasked in healthy subjects [71–75] and animals [76], it is most evident in unmedicated Parkinson’s patients [54], in whom the DA deficiency is profound. This implies a significant asymmetry at baseline: While decreasing the DA levels will have a strong behavioral signature (the central tendency), the effect of increased DA levels will be small (due to a ‘ceiling effect,’ in which the central tendency will continue to be weak). On the other hand, both increases and decreases to the DA level substantially affect the exploration-exploitation trade-off [24, 58–60, 77]. This suggests a more significant central tendency for rewards at baseline, which can be amplified or mitigated by DA manipulations. Below we will find that DA’s effect in ITCs and PDs will depend on its relative contribution to each of the reward estimates and temporal estimates at baseline. Driven by the empirical observations, we take the baseline central tendency to be weaker in the domain of timing than in the domain of rewards.

Finally, it will be useful to distinguish between an animal’s true precision and its estimated precision (or what it perceives its precision to be). True precision refers to the precision with which the animal actually encodes the signal. Estimated precision, on the other hand, determines how heavily to weigh the previously encoded signal against the context, as in Eq. (2). The above treatment assumes perfect calibration between encoding and decoding, so that the animal weighs a signal (during decoding) in perfect accordance with its true encoding precision. However, if the neural substrate of precision is the DA level, then it should be possible to elicit certain biases by selectively manipulating the DA level during decoding. Our main predictions will indeed involve tasks in which the DA level was pharmacologically manipulated after training but immediately before testing.

### Decision making under the Bayesian theory

Having estimated the relevant parameters in the task, how does the animal actually use these parameters to make decisions? Under standard models of action selection, the probability of selecting arm *A_i_* with expected reward 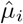 follows a softmax function [78, 79]:

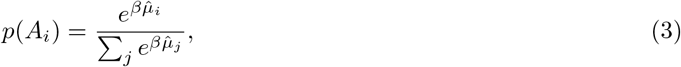

where *β* is the inverse temperature parameter, which controls choice stochasticity. The studies examined in the Results all involve choices between two options; thus, we can restrict our analysis to the case of two arms, *A_l_* and *A_s_*, yielding large and small reward, respectively. Furthermore, each arm not only carries a different reward magnitude but also a different delay period between rewards. Thus, the animal must estimate the arms’ reward *rates* 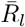 and 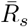 (or ratios of reward magnitude to delay), respectively, in order to maximize its total accumulated reward. Eq. (3) can then be written as

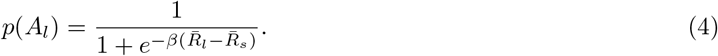

Notice here that the probability of selecting the option yielding the large reward rate depends on the difference between the reward estimates: As the quantity 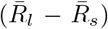 increases, *p*(*A_l_*) increases. Furthermore, by controlling the encoding precisions and thus the central tendencies (either in the temporal or reward domain), DA modulates the estimated difference in posterior means (see horizontal black segments in Fig. 1). A number of authors have argued that DA implements gain control on the values 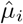 in reinforcement learning tasks, possibly by controlling *β* [60, 66, 69]. The Bayesian theory subsumes the gain control theory by modulating the estimated difference directly.

We have made two important assumptions here. First, our choice rule, though conventional, disregards the contributions of the posterior precisions (i.e., their uncertainties). Recent studies have shown that human behavior in certain bandit tasks is better described by augmented models that incorporate random and directed exploration strategies, both of which make use of the posterior precisions [80–83]. We discuss the augmented model in Supplementary Text 1 and examine its implications for the Bayesian theory.

Our second assumption is about the *shape* of the posterior distributions. As mentioned above, we have assumed Gaussians throughout, with fixed encoding noise. These assumptions are for convenience: Indeed, the uncertainty about an arm should be lower when the arm is more frequently sampled (more well-learned), higher when it is sampled further in the past, and higher for larger-magnitude stimuli [84–87]. Our results will not depend strongly on either the Gaussian assumption or the absolute magnitude of the encoding uncertainty, but only that the central tendency be sufficiently reduced or strengthened under sufficiently high and low precisions, respectively.

## Results

### Dopamine and intertemporal choice

ITCs involve choosing between a small reward delivered soon, and a large reward delivered later. In these tasks, the smaller/sooner delay is held fixed (and is often zero, resulting in immediate reward), while the larger/later delay is varied across blocks. When the delays are equal, animals will overwhelmingly choose the larger option, but as the delay for the larger option gets longer, animals become more and more likely to choose the smaller/sooner option (Fig. 3). This shift toward the smaller/sooner option has traditionally been explained in terms of reward discounting: The promise of a future reward is less valuable than that same reward delivered immediately, and becomes even less valuable as the delay increases. In other words, future rewards are discounted in proportion to the delay required to receive them. Previous computational models have shown this reward discounting to be well-described by a hyperbolic (or quasi-hyperbolic) function [21, 88].

A competing line of thought is that animals seek to maximize their reward rates (or equivalently, the total accumulated rewards in the task) [48, 49, 51], but are limited by a significant underestimation of the post-reward delays in the task [50]. On this view, animals compute the reward rate for each option—i.e., the undiscounted reward magnitude divided by the total trial time—but base the trial time largely on the pre-reward delay. This causes the reward rate for the smaller/sooner option to be disproportionately overestimated compared to that of the larger/later option. This view, much like the discounting view, predicts that animals will choose the larger/later option when its delay is short, but will gradually begin to prefer the smaller/sooner option as the delay is increased. Furthermore, the smaller/sooner option will be preferred in some cases even when it yields a lower reward rate, although this is due to a temporal bias (underestimation of post-reward delays), rather than a choice bias (reward discounting). Note here that we use the term ‘discounting’ to refer to the psychological principle that future rewards are valued less than immediate rewards by virtue of the need to wait for them. Thus, even though in the reward-rate view, rewards are divided by their temporal interval, they are not ‘discounted.’

While the reward-rate interpretation can accommodate the aspects of the data explained by the discounting model, it also captures aspects of animal behavior where the discounting model fails. In particular, Blanchard et al. [50] examined the effect of post-reward delays on behavior. Under the discounting model, behavior depends only on the reward magnitudes and *pre*-reward delays (over which the discounting occurs), and thus should be invariant to changes in the post-reward delays. The authors, however, found that monkeys modified their choices in line with a reward-rate computation, which must take into account both pre- and post-reward delays when computing the total trial time. Interestingly, the best fit to the data required that the post-reward delays be underestimated by about a factor of four, consistent with a bias of timing rather than a bias of choice in explaining animal behavior in ITCs. In what follows, we adopt the reward-rate interpretation in examining DA’s role in ITCs.

Given DA’s effects on reward estimates and durations, it is not surprising that DA would influence behavior in ITCs, where the agent’s task is to maximize the ratio of these two, the reward rate 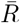:

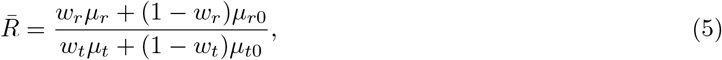

which follows from Eq. (2). Here, 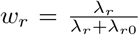, and *μ*_*r*0_, *λ*_*r*0_, *μ_r_*, and *λ_r_* in the numerator represent the prior mean, prior precision, encoding distribution mean, and encoding distribution precision in the domain of rewards, respectively, and similarly for the domain of time in the denominator. Increasing the DA level increases both encoding precisions, *λ_r_* and *λ_t_*.

To understand DA’s overall effect on the ratio 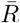, it will be useful to examine manipulations of reward and temporal precision separately. First, let us hold the temporal precisions (and thus the temporal estimates in the denominator) constant. A strong central tendency for the estimated rewards causes an overestimation of the smaller reward and an underestimation of the larger reward, thus promoting selection of the smaller/sooner option compared to baseline. Because increasing DA masks the central tendency, its effect on the reward estimates in the numerator is to promote selecting the larger/later option (Fig. 2A and B, top arrow). Now let us hold the reward precisions constant. In the denominator, a stronger central tendency for the estimated durations causes an overestimation of the sooner duration and an underestimation of the later duration, thus promoting selection of the larger/later option. Because increasing DA masks the central tendency, its effect on the temporal estimates in the denominator is to promote selecting the smaller/sooner option—the opposite of its effect in the numerator (Fig. 2A and B, bottom arrow). Thus the ultimate effect of DA will depend on its relative contribution to the reward and temporal estimates (see Supplementary Text 2 for an analytical derivation).

**Figure 2:**
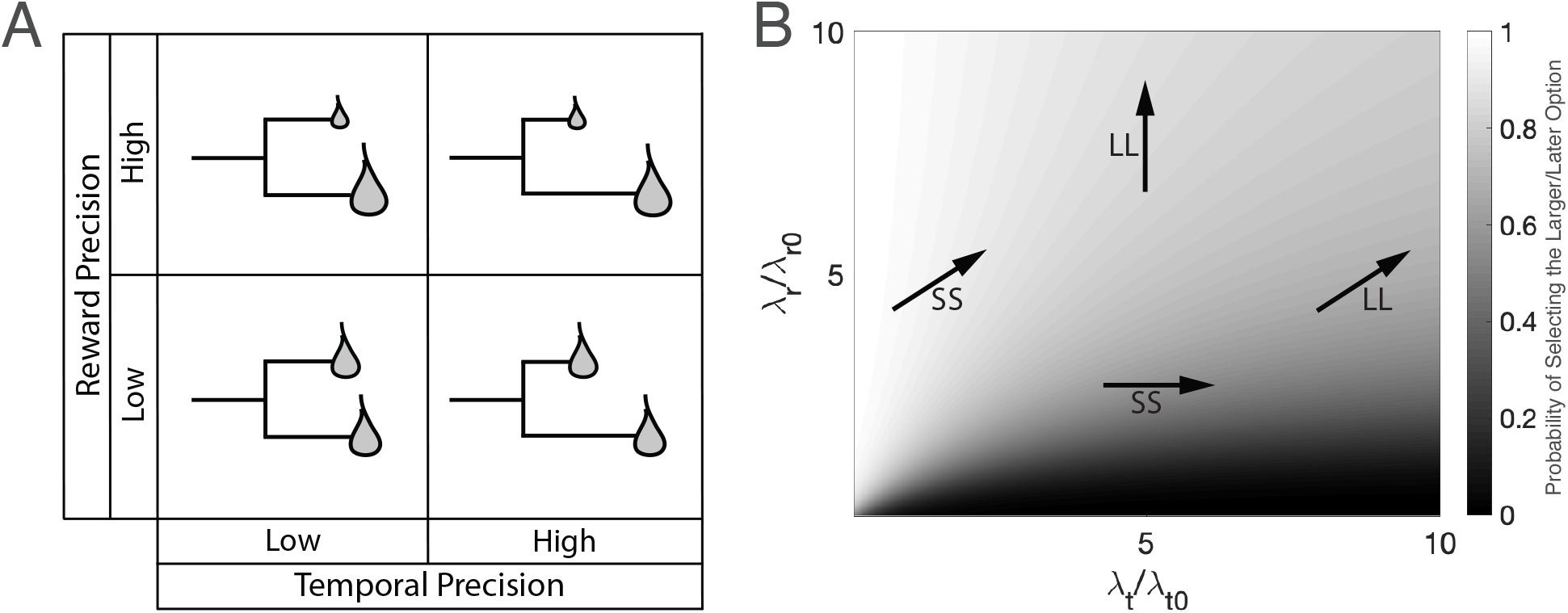
Behavior in ITCs depends on the relative change in reward precision compared to temporal precision. (A) Schematic illustrating the reward and temporal estimates for each of the smaller/sooner and larger/later options under different reward and temporal precisions. Selectively increasing the reward precision (bottom cells to top cells) masks the reward central tendency, making the difference in reward estimates larger. According to Eq. (4), this promotes selection of the larger/later option. On the other hand, selectively increasing the temporal precision (left cells to right cells) masks the temporal central tendency, making the difference in temporal estimates larger. This promotes selection of the smaller/sooner option. (B) Isolines representing pairs of relative precisions that yield the same probability of selecting the larger/later option under Eq. (4). Note that these isolines have different concavities: In the top left, the isolines are concave up (or convex), whereas in the bottom right, the isolines are concave down. Selectively increasing the reward precision promotes the larger/later option (top arrow), whereas selectively increasing the temporal precision promotes the smaller/sooner option (bottom arrow). Based on empirical findings, we assume that the temporal precision at baseline is high, compared to the baseline reward precision (each normalized by its prior precision). This means that DA’s net effect is to promote the larger/later option (right arrow). If, however, the temporal precision is sufficiently reduced, DA’s net effect will be to promote the smaller/sooner option (left arrow). Plotted on each axis is the ratio of encoding and prior precisions, which determines the central tendency: 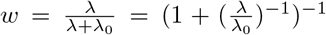. For illustration, we have chosen *μ_r_* = 1 and 4, and *μ_t_* = 2 and 6, for the smaller/sooner and larger/later options, respectively, and *β* = 10. LL: increase in probability of selecting the larger/later option, SS: increase in probability of selecting the smaller/sooner option, *λ_t_*: temporal encoding precision, *λ*_*t*0_: temporal prior precision, *λ_r_*: reward encoding precision, *λ*_*r*0_: reward prior precision.

As discussed in the previous section, the central tendency at baseline DA levels is stronger for reward estimates than temporal estimates. It follows that the central tendency in the numerator dominates DA’s influence in ITCs (Fig. 2B, right arrow). Under normal conditions, then, the framework predicts that increasing DA will promote the larger/later option, or behavior consistent with lower impulsivity under higher DA levels (Fig. 3E).

**Figure 3:**
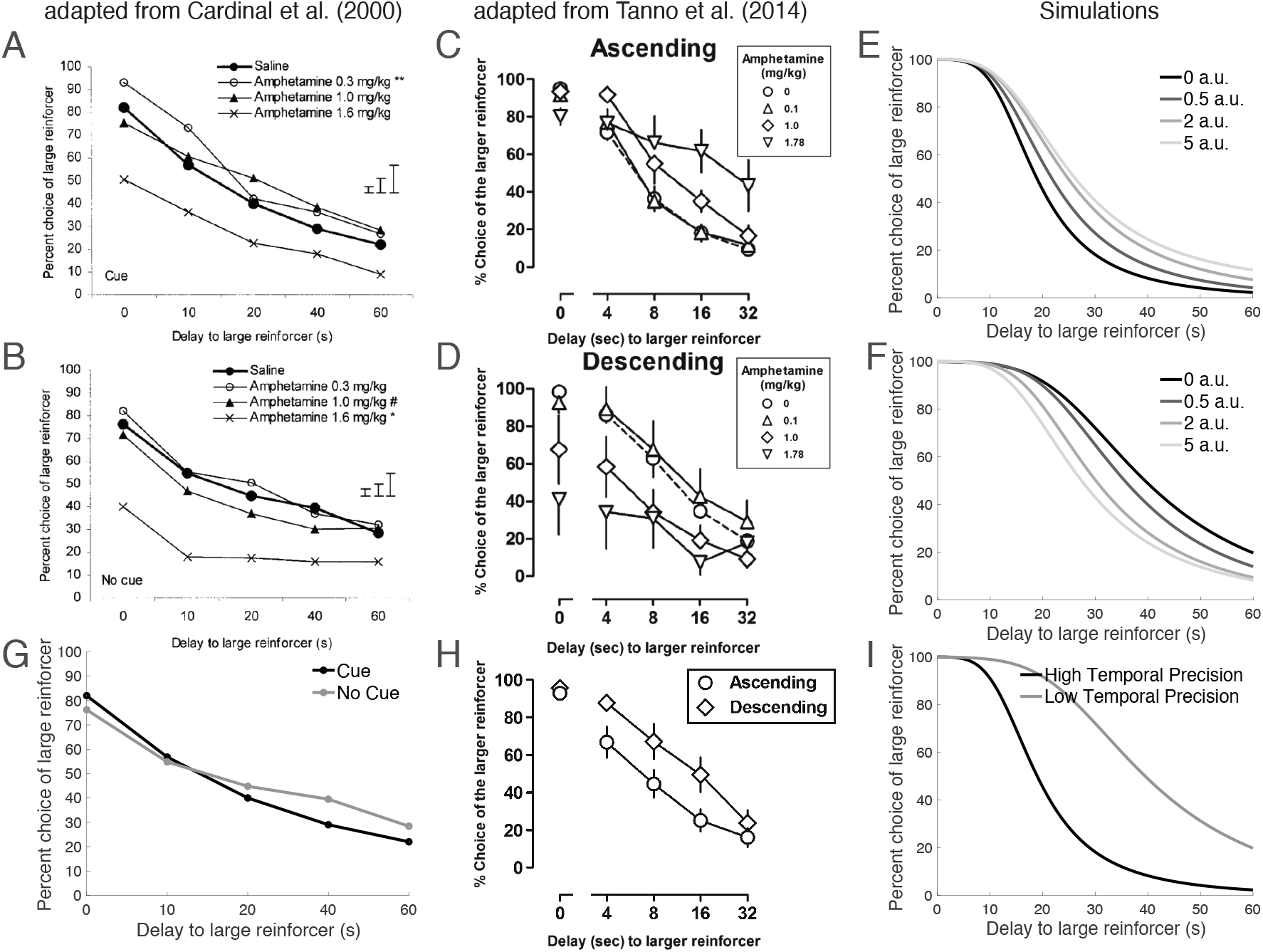
DA agonists promote selection of the larger/later option when the temporal precision is high and the smaller/sooner option when the temporal precision is low. (A) Cardinal et al. [16] trained rats on an ITC in which the animals must choose between a reward of magnitude 1 delivered immediately and a reward of magnitude 4 delivered after a delay that varied across blocks. After training, the authors administered DA agonists and examined changes in the animals’ behaviors. When a cue was present during the delay period, the authors found that the animals seemed less impulsive under DA agonists, or discounted future rewards less. The 0.3 mg/kg dose, but not the other doses, reached statistical significance (\*\**p* < 0.01, main effect of DA agonist). (B) However, when a cue was absent during the delay period, the animals appeared more impulsive with higher doses, i.e., discounted future rewards more strongly (\**p* < 0.05, main effect of DA agonist; ^#^*p* < 0.05, agonist-delay interaction). For (A, B), vertical bars denote the standard error of the difference between means for 0.3, 1.0, and 1.6 mg/kg relative to saline, from left to right. (C) Tanno et al. [33] administered a similar task, but varied the order in which the delays were presented. When the delays were presented in an ascending order, the rats seemed less impulsive with higher doses of DA agonists. (D) However, when the delays were presented in a descending order, the rats seemed more impulsive with higher doses. (E) Our model recapitulates these effects: Under high temporal precision, such as in the presence of a visual cue during the delay (cue condition) or as suggested empirically by measuring response variability (ascending condition; Supplementary Text 3), DA’s effect on the reward estimates will dominate in ITCs, which promotes selection of the larger/later option. (F) On the other hand, under sufficiently low temporal precision, DA’s effect on the temporal estimates will dominate, which promotes selection of the smaller/sooner option. (G) At baseline, responses in the no-cue condition are biased toward the larger/later option compared to the cue condition. Note that any zero-delay difference cannot be due to a difference in the cues, since the tasks are identical in the absence of a delay. It is not clear whether these differences are statistically significant, as error bars were not provided for the saline conditions (although when the conditions were tested immediately before drug administration began, the difference was not statistically significant). Panel reproduced from the saline conditions in (A) and (B). (H) Similarly, at baseline, responses in the descending condition are biased toward the larger/later option compared to the ascending condition. (I) Our model recapitulates these effects: Selective decreases to the temporal precision promote the larger/later option. For (E, F, I), see Supplementary Text 3 for simulation details. a.u.: arbitrary units of DA.

This prediction matches well with empirical findings, as the majority of studies have found administering DA agonists to decrease impulsivity in ITCs [15, 22–28] (see [29] for a recent review). For instance, Cardinal et al. [16] trained rats on an ITC involving a small reward delivered immediately and a large reward delivered after a delay that varied across blocks. After training, the authors administered DA agonists and tested the animals on the task. While the effect is smaller than in other studies (e.g., compare with Fig. 3C), the authors found that the DA agonists promoted selection of the larger/later option when a visual cue was present throughout the trial (Fig. 3A).

This prediction is based on the empirically motivated result that DA’s effect on the reward estimate dominates its overall effect in ITCs. However, it should be possible to elicit exactly the opposite result—an increased preference for the smaller/sooner option with DA—under conditions where the central tendency of temporal estimates dominates. For instance, timing precision is affected by the inclusion of temporally-informative cues [84, 89, 90] as well as manipulations of the interval salience, presumably due to changes in the animal’s alertness [91]. Then removing these cues and selectively decreasing the salience during the delay period should promote the temporal central tendency and, if significant enough, overwhelm the central tendency of rewards in the numerator (Fig. 2, left arrow). Cardinal et al. [16] examined exactly this manipulation: The authors found that DA, on average, promoted selection of the larger/later option only when a salient, temporallyinformative visual cue was selectively available during the delay period. Otherwise, DA uncharacteristically had the opposite effect (Fig. 3B), as predicted when the temporal precision is sufficiently reduced (Fig. 3F).

It is important to note that DA manipulations can mediate preference reversals, which is captured by our model. For example, for the 8- and 16-second delays in Fig. 3C, the animal at baseline prefers the smaller/sooner option (chosen more than 50% of the time). But with high enough doses of DA agonists, it eventually comes to prefer the larger/later option. This empirical finding is important because it rules out hypotheses in which DA simply amplifies or mitigates existing preferences. For instance, and as mentioned above, a number of authors have proposed that DA serves a ‘gain control’ function on the action values during decision making [67–70]. This would predict that preferences should become more extreme with higher DA levels: Preferences above the indifference (50%) line should increase, and those below the indifference line should decrease, which is inconsistent with the empirical results.

Though the majority of studies have found behaviors consistent with a negative correlation between impulsivity and DA, Cardinal et al. [16] found the opposite effect when the cue was selectively absent during the delay period, and we showed that the Bayesian framework captures this effect. We are aware of one other manipulation that may cause this opposite effect: In tasks where animals are trained on different delays for the larger/later option, Tanno et al. [33] have reported that DA’s effect depends on the ordering of the delays. In particular, they found that DA agonists seemed to promote choosing the larger/later option, in line with most other studies, when the delays were presented in an ascending order. However, if the delays were presented in a descending order, DA agonists had the opposite effect (see also [92]). This finding would be consistent with our framework, if the temporal precision in the ascending case were higher than that in the descending case (Fig. 3C,D). This may indeed be the case, as when learned in an ascending order, the animals’ temporal behavior (i.e., the timing of the animal’s lever press) was less variable than when learned in a descending order (Supplementary Text 3). An important limitation here is that this result does not control for the animal’s motivational state: It is possible that the ordering effect influences not the animal’s temporal precision but its motivation, leading to less temporally precise behavior. It is not clear *why* such an ordering effect exists, although one possibility is that this arises from a primacy effect in the inference about the temporal sequence [93–96], as the initial temporal precision is higher for the short delays due to Weber’s law [84, 89, 90], and potentially also due to the incentive structure (the animal is more incentivized to attend to blocks with higher reward rates, i.e., those with short delays) [57, 97].

Third, the Bayesian framework makes a counterintuitive prediction about the relationship between baseline performance in ITCs and the effect of DA. According to our model, selectively increasing the temporal precision promotes the smaller/sooner option. However, DA’s effect, when the temporal precision is already high, is to promote the *larger/later* option (compare bottom and right arrows in Fig. 2). This implies that conditions in which DA agonists promote the larger/later option will be conditions in which animals are, at baseline, more likely to select the smaller/sooner option. The authors of both studies above indeed observed this relationship: For both the cue and ascending conditions, animals were more likely to select the smaller/sooner option at baseline, compared to the no-cue and descending conditions, respectively (Fig. 3G,H), as predicted (Fig. 3I). Note, however, that this effect may also be due to baseline differences in the speed of the ‘internal clock’: There is some evidence to suggest that slower clocks are correlated with lower temporal precision [57]. This means that, in tasks with low temporal precision, the animal may perceive the interval to be shorter than in tasks with high temporal precision, which may make it more appealing (shorter intervals result in larger reward rates). We examine this point at length in Supplementary Text 4. Interestingly, combined with the correlation between clock speed and temporal precision, the Bayesian theory makes another prediction: that temporal intervals that are underestimated (slow internal clock) should not be well-learned (low encoding precision, resulting in strongly overlapping posteriors due to the central tendency). As mentioned previously, Blanchard et al. [50] indeed observed that post-reward delays are underestimated and not well-learned, but that this learning can be salvaged if the temporal intervals are made salient, as our model predicts (Supplementary Text 4).

It should be noted that, while our model is concerned with the main effect of DA manipulations, animal response profiles seem also to profoundly diverge in the descending task for the smaller delays (note splaying of response profiles in Fig. 3D). Our model can accommodate this result: Because of noisy learning, the encoding of current estimates will be biased toward estimates from previous blocks, a form of ‘within-arm’ contextual influence. This effect should be more apparent under higher DA levels, because of a silencing of the central tendency. We expand on this point in Supplementary Text 5.

Finally, having examined DA’s effects on behaviors in interval timing and measures of impulsivity, we can also examine how the behavioral phenotypes covary with each other. Our model predicts that—due to natural differences in DA levels within a species—animals that are more precise timers should also appear less impulsive in ITCs, as has indeed been observed [98, 99] (Supplementary Text 6).

We have sought to highlight here a link between temporal precision and the effect of DA in ITCs. Under the Bayesian theory, reward and temporal estimates normally regress to their contextual means in inverse proportion to the encoding precisions, and increasing the DA level mitigates this regression. This increases both the estimated cost (delay) and benefit (reward) of the larger/later option. When the temporal precision is already high (negligible regression to the temporal mean), the increase in benefit dominates DA’s effect, and the animal becomes more likely to select the larger/later option. When the temporal precision is sufficiently low (strong regression), the increase in cost dominates, and the animal shifts its preference toward the smaller/sooner option. Note here that our focus on temporal, rather than reward, precision is driven by the experimental paradigm: Of the four estimated parameters—small reward, short duration, large reward, and long duration—only the long duration is varied across blocks, so it is not surprising that variations of the ITC would be characterized by different temporal precisions. In principle, a similar analysis can be conducted for manipulations of the reward precisions.

### Dopamine and probability discounting

We now extend the Bayesian analysis to risk-seeking and PDs. We follow a similar outline to the previous section in showing that, like for the case of impulsivity and ITCs, the findings of DA in PDs are fully captured by the Bayesian theory, and do not necessarily reflect changes in risk preferences at all.

St Onge and Floresco [46] tested rats on a task in which they had to choose between an arm yielding 1 pellet with 100% probability (the small/safe option) and another arm yielding 4 pellets but with a probability that was varied across blocks (the large/risky option). As the probability decreased, the rats became less likely to choose the large/risky option. After the rats achieved stable performance in each block, the DA agonist amphetamine was administered. The authors found that the DA agonist induced a tendency to select the large/risky option, which was taken to suggest that increasing the DA levels may promote risk-seeking behaviors. However, in a follow-up study [47], the authors found that the ordering of the blocks mattered: The DA agonist only promoted the large/risky option when the probability of reward decreased with blocks. When the order was reversed, the DA agonist had the opposite effect (Fig. 4A,B).

**Figure 4:**
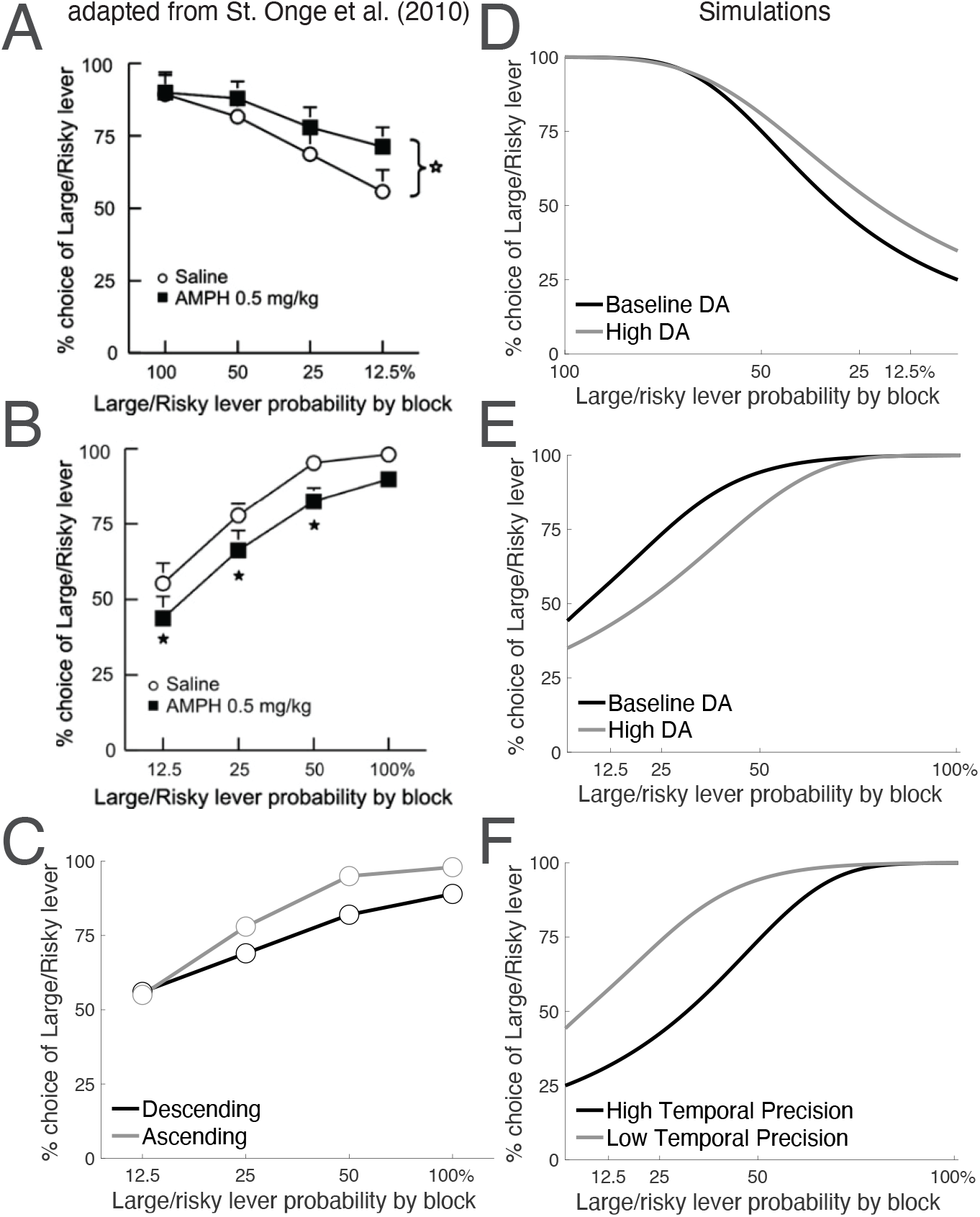
DA agonists promote selection of the large/risky option when the temporal precision is high and the small/safe option when the temporal precision is low. (A) St Onge et al. [47] trained rats on a PD in which the animals chose between 1 pellet delivered on 100% of trials and 4 pellets delivered with some probability that varied across blocks. After training, the authors administered the DA agonist amphetamine. They found that, when the probabilities were presented in a descending order, amphetamine induced an increase in the tendency to select the large/risky option. (B) However, when the probabilities were presented in an ascending order, amphetamine had the opposite effect—inducing a decrease in the tendency to select the large/risky option. (C) At baseline, responses in the ascending condition are biased toward the large/risky option compared to the descending condition. Panel reproduced from the saline conditions in (A) and (B). As per Fig. 3G, it is unclear for which doses the differences in choice behavior are statistically significant, although visual inspection suggests a main effect of block order. (D) Our model recapitulates these effects: Under high temporal precision, DA’s effect on the reward estimates will dominate, which promotes selection of the large/risky option. (E) On the other hand, under sufficiently low temporal precision, DA’s effect on the temporal estimates will dominate, which promotes selection of the small/safe option. (F) Selective decreases to the temporal precision promote the large/risky option. See Supplementary Text 3 for simulation details.

The Bayesian theory predicts this finding. At a conceptual level, this is because the task sets up a trade-off between two attributes (reward magnitude and risk), whose central tendencies push the reward rate in opposite directions: An animal that learns the reward magnitudes well and largely disregards the risks will view the large/risky option as superior, whereas an animal that learns the risks (probabilities) well but disregards the reward magnitudes will prefer the small/safe option.

To evaluate the theory’s predictions more concretely, we note that there are a number of ways for an animal to learn the reward rate, and therefore a number of ways to set up the two central tendencies. For instance, the animal may estimate the reward magnitude and reward probability separately and take their product. Alternatively, the animal may estimate the reward magnitude and average delay time between two rewards for each option, and take their ratio, as in Eq. (5). While both approaches can accommodate the empirical results, we will assume the latter for two reasons: First, this approach allows for a direct comparison with ITCs, as delay discounting and probability discounting elicit similar behaviors [100, 101], similar types of intolerance [102, 103], and a common neural substrate [44, 104]. Second, after Tanno et al. [33], if the animal is indeed computing the ratio of reward magnitude to the delay, then the block order manipulation makes a clear prediction about the temporal precision: A block order involving sequentially longer delays (here, the descending condition) results in higher temporal precision than an order involving sequentially shorter delays (here, the ascending condition). As was the case for the ITC, this will be the key variable determining DA’s overall effect.

Thus, to maximize its reward rate, we assume an animal estimates the reward magnitude and average delay between rewards for each option, and computes their ratio. For the reward magnitudes, the central tendency promotes selection of the small/safe option. Increasing DA silences this effect, and thus promotes selection of the large/risky option. On the other hand, for the temporal interval, the central tendency promotes selection of the large/risky option, which involves a longer waiting time between rewards. Increasing DA therefore promotes selection of the small/safe option here—the opposite of its effect for reward magnitudes. Thus, once again, DA’s overall effect depends on its relative contribution to each term. Following the previous section, the temporal precision is predicted to be low in the ascending condition (which involves sequentially shorter delays) compared to the descending condition (which involves sequentially longer delays). It follows that the temporal central tendency dominates in the ascending condition and the reward central tendency dominates in the descending condition, thus predicting the empirical findings (Fig. 4D,E).

Our theory also predicts that selectively increasing the temporal precision, without manipulating DA, should also make animals more likely to select the small/safe option. Indeed, by examining the baseline (saline) task for each of the ascending and descending conditions, we find that animals were more likely to select the small/safe option in the descending condition at baseline compared to the ascending condition (Fig. 4C), as predicted (Fig. 4F).

Finally, having examined temporal and probability discounting separately, we briefly mention the ‘rat gambling task,’ a four-armed bandit task in which both the delay periods and reward probabilities (in addition to the reward magnitudes) are varied across arms. A well-replicated finding in this task has been that the arm yielding the largest reward magnitude is the second most frequently selected arm, even though it yields the lowest reward rate. In addition, amphetamine administration tends to disrupt selection of the arm yielding the highest reward rate in favor of that yielding the second highest reward rate [105]. The Bayesian theory can accommodate both findings (Supplementary Text 7).

## Discussion

We have shown here that DA’s effects in ITCs and PDs are well-described by a Bayesian framework in which animals maximize their reward rates. Under this view, DA controls the relative influence of context in computing the reward and temporal estimates, whose ratio forms the reward rate. Notably, the discounting-free model successfully predicts that DA agonists should promote selection of the larger/later and large/risky options under high temporal precision, but should have exactly the opposite effects when the temporal precision is sufficiently low. The Bayesian view thus provides a principled framework for why DA would appear to inhibit impulsive and risky choices in some paradigms but promote them in others.

We have followed previous theoretical and experimental work in adopting a discounting-free model of choice behavior. However, our results do not necessarily rule out temporal or probability discounting more generally, nor a role for DA in these processes. For instance, and as mentioned in the Introduction, humans tend to prefer smaller/sooner options even in the absence of repeated trials that make reward-rate computations meaningful. But why discount future rewards in the first place? One influential hypothesis from economics is that future rewards are discounted because of the risks involved in the delay [106]. For example, a competitor may reach the reward first, or a predator may interfere in the animal’s plans to collect the reward. As the delay increases, these alternative events become more likely, and the expected reward (the average over all alternatives) decreases. Another idea is that subjects respond *as if* they will have repeated opportunities to engage in the same task [107], thus mimicking the reinforcement learning problem that defines the animal variant of ITCs. More recently, Gabaix and Laibson [108] have argued that reward discounting may be due to the simulation noise involved in mentally projecting into the future: With later rewards, subjects must mentally simulate further into the future, so the simulation noise increases, and the precision decreases. Assuming a Bayesian framework with a prior centered at zero, the reward estimates will be closer to zero when rewards are more distant in the future, i.e., rewards are discounted with time (see also [109] for an extension of this hypothesis).

Interestingly, as mentioned in the Introduction, DA seems to have the opposite effect in the human variant of the task than in the majority of animal experiments, with a promotion of the smaller/sooner option with higher DA levels. That DA may serve a qualitatively different function in the human variant is not completely unexpected, given the substantial differences in the experimental paradigms. Notably, in the human variant, (1) the subject does not actually experience the pre-reward delay, (2) there is no post-reward delay, (3) the subject does not necessarily receive an actual reward, (4) the subject may experience a single trial of this task, whereas animals are trained on many trials, and (5) the hypothetical delay is on the order of days (or months) and not seconds. Experience and repetitions may prove critical for our reinforcement learning task, and delays on the order of days engage different timing mechanisms than those on the order of seconds-to-minutes [110], which is the duration over which DA’s central tendency effect has been observed. Nonetheless, the human findings may still be reconcilable with our framework under the ‘repeated opportunities’ hypothesis of Myerson and Green [107] mentioned above: It is possible that the temporal uncertainty surrounding durations that are not experienced, and that are on the order of days, is large and thereby dominates DA’s central tendency effects. Thus DA agonists would be predicted to promote the smaller/sooner option.

Our framework leaves open a number of theoretical and empirical questions. First, our model takes DA to control the encoding precision, a property inherited from the Bayesian timing model of DA and further motivated by theories of DA as overcoming the cost of attention [57, 97]. However, our results only require that DA control the ratio of the encoding precision to the prior precision but not necessarily the encoding precision itself. Instead, it is certainly possible that increasing DA decreases the prior precision, as some authors have proposed [53]. Interestingly, this ambiguity is not specific to theories of DA, and has been a point of debate for some Bayesian theories of autism as well (compare weak priors [111] with strong likelihoods [112]).

A second open question concerns our assumption that estimates of the reward magnitude are biased by a central tendency effect. Thus far, this has been inferred mainly from exploration-exploitation paradigms (see [64] for a more direct examination), but a dopaminergic modulation of reward estimates has not, to our knowledge, been observed directly. Driven by the experimental literature, we have therefore focused our simulations on manipulations of *temporal* precision. Our work then opens the door to a fruitful line of experiments with novel predictions: For instance, one can develop ITCs and PDs where the large reward is varied rather than the delay or risk. Our framework predicts that DA agonists will promote the larger/later and large/risky options only when *reward* precision is low at baseline, and the smaller/sooner and small/safe options when reward precision is high. On the other hand, selectively increasing the reward precision will always promote the larger/later and large/risky options (Fig. 2). Thus, once again, by simply controlling the central tendency, DA agonists will appear to inhibit impulsivity and risk-seeking under some conditions, but promote them in others.

Third, having adopted an algorithmic view of DA’s function, it remains for future work to ask how the Bayesian theory is actually implemented neurobiologically. Notably, DA exerts different effects depending on the postsynaptic receptor subtype: In reinforcement learning studies, midbrain DA neurons project to the striatum onto neurons primarily expressing either D1 or D2 receptors, which segregate anatomically into largely separate basal ganglia pathways [113] and seem to serve opposite purposes [114, 115], both in their fast-[116] and slow-timescale [69, 117, 118] activities. Asymmetries in receptor-mediated effects extend into interval timing studies (compare D1-mediated [119–123] with D2-mediated [119, 120] effects), and DA’s effects also depend on enzymatic activity [124, 125] and projection site [126, 127]. Bridging the algorithmic and implementational levels of the Bayesian theory will be a necessary next step toward a more complete theory of DA.

Finally, we have examined in this work how manipulations of DA affect behavior in the ITC and PD, but it is interesting to ask what variables determine the DA level in the first place. An influential proposal has been that the tonic DA level is set by the average reward availability in an environment [128], as has also been suggested empirically [129]. One unifying interpretation of the average reward and Bayesian theories is that, in high reward-rate environments, animals are more incentivized to attend to a task, and thus encode the relevant parameters with higher precision. In this manner, DA connects the encoding stage (learning the parameters) with the decoding stage (combining the learned parameters with contextual information, the focus of this paper) [57]. It will remain for future work to build upon and experimentally validate coherent theories of DA within an encoding-decoding framework.

To our knowledge, this is the first framework that can accommodate the seemingly conflicting effects of DA in measures of impulsive choice and risk-seeking across experimental conditions. Nonetheless, our aim throughout this work is not to rule out a role for DA in true impulsivity and risk-seeking, but rather to show how a single Bayesian framework can accommodate a wide range of otherwise perplexing behavioral and pharmacological phenomena.

## Acknowledgements

The authors are grateful to Rahul Bhui for comments on an earlier draft of the paper.

## Author contributions

J.G.M. and S.J.G. developed the model and contributed to the writing of the paper. J.G.M. analyzed and simulated the model, made the figures, and wrote the first draft.

## Institutional review board

This is a theoretical study which does not describe any new data.

## Data and code availability

Source code for all simulations can be found at www.github.com/jgmikhael/bayesiantheory.

## Supplementary Information

### 1 Random and directed exploration

We have assumed that animals make decisions based on the softmax choice rule with a fixed inverse temperature parameter *β*, taking as inputs the estimated reward rates 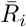. Though widely used, this model may be simplistic in that it does not take the animal’s uncertainty into account. For instance, more uncertain contexts should elicit more exploratory behaviors [80]. In line with this intuition are two influential models of uncertainty-based action selection: one in which randomness is added to choice behavior, and one in which choices are directed toward more uncertain options. Here we consider the effect of each exploration strategy on our main predictions.

One classical random exploration strategy, known as Thompson sampling [130], is to make choices more stochastic in proportion to the total uncertainty in the environment. Under Thompson sampling, the animal draws one sample from each option’s posterior over reward rates and chooses the option with the largest sampled reward rate. Using the notation in the main text, the animal draws samples 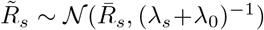 and 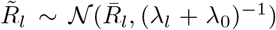 from options *A_s_* and *A_l_* respectively, and chooses option *A_l_* if and only if 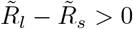. This occurs with probability

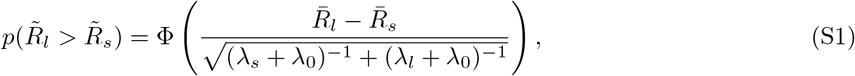

where Φ(·) is the cumulative distribution function of the standard normal distribution. When the encoding precisions are very large, the denominator on the right-hand side approaches 0, and the probability approaches 1. Intuitively, when the overlap between the two posterior distributions is minimal, the sample drawn from *A_l_* will always be greater than that drawn from *A_s_*, and as the overlap increases, the probability of selecting *A_s_* increases.

Note here that the denominator on the right-hand side can be thought of as the temperature parameter *β*^−1^, which is a function of the total uncertainty in the environment: When the total uncertainty is large, *β*^−1^ is large, and choices are highly stochastic. As the total uncertainty decreases (e.g., by sampling, and learning about, the environment), *β*^−1^ decreases, and choices become more deterministic. Under the Bayesian theory of DA, increasing DA increases *λ_s_* and *λ_l_*, thereby decreasing *β*^−1^. Thus DA’s effect on the denominator is to make choices more deterministic. This is reminiscent of the gain control view of DA discussed in the main text.

A second exploratory strategy is directed exploration. Here, an option’s uncertainty functions as a bonus that is added to its estimate, thus promoting exploration of the more uncertain option. In keeping with the softmax function, we can write:

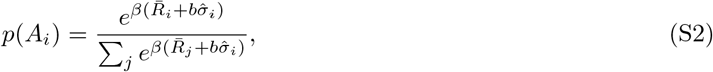

where 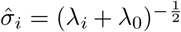 is the posterior standard deviation and *b* > 0 is the bonus coefficient which controls the contribution of uncertainty in making decisions. The uncertainty term 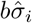 benefits options with higher uncertainty.

Though the empirical evidence is limited in animals, previous work has shown that humans use all three strategies in standard bandit tasks [83]. Gershman [83] modeled choice behavior as a combination of the three strategies,

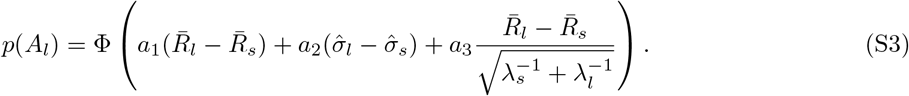

The first, second, and third terms in the argument on the right-hand side represent cases where choice behavior depends on the difference in estimated reward rates, the difference in posterior standard deviations, and the difference in samples drawn from each posterior distribution, respectively. Gershman [83] found that this augmented choice rule afforded the best quantitative account of human behavior in the bandit task, with *a*_1_ > *a*_3_ > *a*_2_ > 0. (Note, however, that contextual priors were not incorporated into this analysis.) In general, our results hold when the standard softmax function assumed in the main text dominates the augmented choice rule. When the directed exploration strategy dominates, our results furthermore hold when 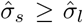, because increasing DA (decreasing 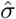) more strongly reduces 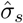, so the difference 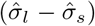 increases. It is reasonable to assume that 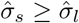, as animals will predominantly sample the more rewarding arm, thus reducing their uncertainty about it.

### 2 Dopamine’s effect depends on its relative contribution to reward vs. temporal estimates

Our results in the main text rest on the intuition that in reward-rate computations, DA’s effect on *reward* estimation will dominate when temporal precision is high, but its effect on *temporal* estimation will dominate when temporal precision is low. We show this analytically here.

As increasing the DA level increases the encoding precisions, we begin by taking the derivative of 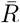 in Eq. (5) with respect to the DA level *d*:

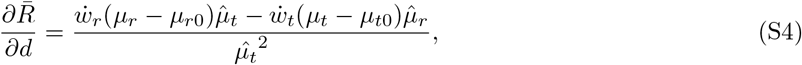

where 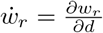 and 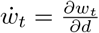. We are interested in how DA affects 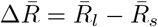, the difference between the larger/later reward rate estimate 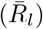 and the smaller/sooner reward rate estimate 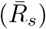. According to the choice rule in Eq. (4), this quantity determines the animal’s behavior.

When the temporal precision is sufficiently high, 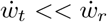. Intuitively, 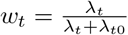 approaches 1, so small changes in DA do not affect it very strongly, compared to *w*. Formally, 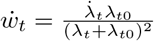 and 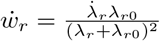, where 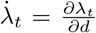 and 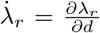. Because the prior precisions are finite, we require that 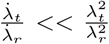, so that 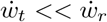.

It follows that, in Eq. (S4), the first term in the numerator dominates:

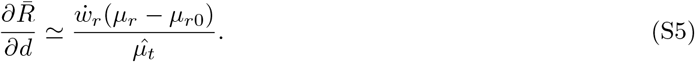

The term in the parentheses is positive for the larger/later option and negative for the smaller/sooner option. Then, 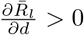 and 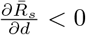. It follows that 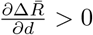, so increasing DA promotes the larger/later option.

Similarly, when the temporal precision is sufficiently low, 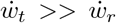. Formally, we require that 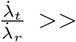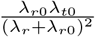, so that small changes in DA strongly affect *λ_t_*, and, by extension, *w_t_*. In this case, the second term in the numerator of Eq. (S4) dominates:

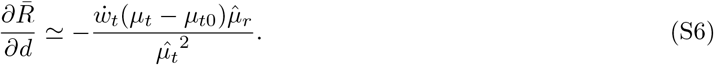

The term in the parentheses is positive for the larger/later option and negative for the smaller/sooner option. Then, 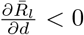 and 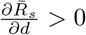. It follows that 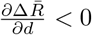, so increasing DA promotes the smaller/sooner option.

Note here that the approximation in Eq. (S6) depends on the durations being different for each option. Otherwise, (*μ_t_* − *μ_t_*_0_) = 0, and the first term in the numerator in Eq. (S4) will always dominate, regardless of how low the temporal precision is. In this case, increasing DA will always promote the larger option. Said differently, if the delays are equal, the task reduces to a simple two-armed bandit task (the options are equivalent except for a difference in reward magnitudes), and our framework predicts that increasing DA will always promote the larger option.

### 3 Simulation details

For our results to hold, we require that the mapping between DA and the encoding precision be monotonic over the relevant domain. It will be convenient to refer to the encoding standard deviation 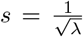. We arbitrarily set 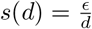, where *ϵ* represents signal-independent noise, and *d* is the DA level. We treat a DA level of 1 as the baseline (e.g., saline) condition. Increasing the DA level *d* decreases the standard deviation *s* and thus increases the precision *λ*, as required.

We assume, as in the Results, that the central tendency is more profound in the domain of rewards than in the domain of timing under normal conditions (high temporal precision). For conditions in which the temporal precision is selectively and sufficiently reduced, we increase the encoding standard deviation by a factor of *g*.

The prior mean and standard deviation were set to the mean and standard deviation of the distribution of stimuli. The prior precision *λ*_0_ is an inverse function of the prior variance 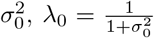, where the added ‘1’ in the denominator is so the precision does not go to infinity when the two stimuli are equal (a form of Laplace smoothing).

For Fig. 3, we set the reward magnitude for the smaller/sooner and larger/later options to 1 and 4, respectively. The total trial length was 100 seconds, and the post-reward delay was 100 minus the pre-reward delay. We assumed the post-reward delays were underestimated by a factor of 10. We set *ϵ* to 0.5 and 4 for rewards and timing, respectively, and *g* to 5. As per the Results, we assumed that the encoding standard deviation increases by the same factor (here, 10), compared to the encoding standard deviation for pre-reward delays. Finally, the agent makes decisions based on the softmax choice rule in Eq. (3) with *β* = 30.

For Fig. 4, we set the reward magnitude for the small/safe and large/risky options to 1 and 10, respectively. The true average delay between two rewards is the product of the total trial length and the average number of trials between two rewards (or the inverse of the block probability, e.g., when reward probability is 12.5%, the average number of trials between two rewards for the large/risky option is 1/0.125 = 8). The animal’s estimate of the average delay between two rewards is the sum of the pre-reward delay and the post-reward delay. The total trial length was 40 seconds, and the pre-reward delay for a single trial was 7 seconds. The post-reward delay was the average duration between rewards minus the total pre-reward delay across trials. We assumed the post-reward delays were underestimated by a factor of 10. We set *ϵ* to 2 and 7 for rewards and timing, respectively, and *g* to 10. We increased *d* by 10 to capture the effect of DA agonists. Finally, the agent makes decisions based on the softmax choice rule in Eq. (3) with *β* = 10.

For Fig. S1 (see Supplementary Text 5 for description), we set *α* = 0.2 across 5 blocks, reward magnitudes of 1 and 3 for the small and large options, respectively, and standard deviations of 0.5 and 2 for the reward and temporal estimates, respectively. Trial length was set to 50, *g* = 10, *β* = 20, and the effect of DA is as above. We arbitrarily set the block-history precision to be equal to the encoding precision. For illustration, we’ve increased the number of blocks to 100, and therefore set 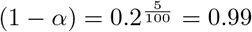.

For Fig. S2, the DA level was varied between 0.5 and 1.5 to mimic natural differences across animals, while being centered at 1. DA levels were sampled logarithmically between these two extremes. For the ITC, the reward magnitudes were 1 and 3, and the post-reward delay was 120 seconds. The short duration was 2.5, 5, 10, or 30 seconds, and the long duration was always 30 seconds. The mean log odds were computed by averaging over the log odds for each temporal pair. We assumed the post-reward delay was underestimated by a factor of 10. We set *E* to 0.8 and 2.7 for rewards and timing, respectively. An inverse temperature parameter of *β* = 180 was required to match the data well, although the mismatch between this *β* value and that of the previous experiments may in part be due to the arbitrarily defined effect of DA on the encoding precision (in Eq. (4), *β* is multiplied by the posterior mean difference, whose relationship with DA is monotonic, but arbitrarily set). All other parameters are identical to those used in the experiments above. Finally, for the bisection task, the probability of selecting the ‘long’ response was the probability of the long duration for each time point, divided by the sum of probabilities of the short and long durations. The stochasticity parameter *σ* was fit to the softmax function in Eq. (3), where *σ* = *β*^−1^.

For Fig. S3, the advantageous and disadvantageous pairs each constitute a context, and the full distribution of four stimuli constitutes the hyper-context. The estimated mean is then:

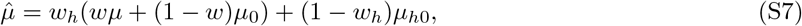

where 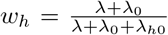, *μ*_*h*0_ is the hyperprior mean, and *λ_h_*_0_ is the hyperprior precision. The agent makes decisions based on the softmax function in Eq. (3) with *β* = 40. To illustrate that the Bayesian theory can capture the results over a restricted parameter space, we arbitrarily set the reward precisions to be large compared to the temporal precisions, and the precisions for the advantageous pair to be large compared to those of the disadvantageous pair. We set the encoding standard deviations to be 0.01 and 1 and the temporal encoding standard deviations to be 2 and 11, for the advantageous and disadvantageous pairs, respectively. The choice latency was set to 2 seconds, the inter-trial interval was 5 seconds, and the collection latency was set to 5 seconds.

Finally, for Tanno et al. [33], the assertion of a difference in temporal precision between the ascending and descending conditions is based on Table 2 in [33]. The authors report the animals’ timing of the lever press for the saline conditions as 1.6 s for both ascending and descending conditions, with standard errors of the mean (SEM) of 0.1 and 0.3, respectively. Both groups consisted of 6 animals; thus, the standard deviation is 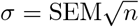, or 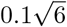 and 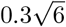, respectively. To test for equality of variances for 2 groups with 6 animals each, the degrees of freedom are 1 and 5, so *F*_1,5_ = 0.11 and *p* = 0.03 < 0.05. Thus we can reject the null hypothesis that the two precisions are equal.

### 4 Clock speed and temporal precision

There is an interesting coupling in the interval timing literature wherein increasing DA both increases the speed of the internal clock [131–135] and masks the central tendency effect. This relationship may be causal, as clock speed may be the mechanism through which precision is modulated (e.g., see [57]). Furthermore, previous theoretical work has argued that precision will only increase when properly incentivized [109, 136–138]—i.e., when an increase in precision improves performance. This would imply that the clock should slow down in tasks in which precision does not improve performance as well as during post-reward delays or intertrial intervals, when the animal has less control over the outcomes. These predictions have some empirical support [50, 139]. Normative arguments notwithstanding, in this section it will suffice to treat the coupling between clock speed and temporal precision as an empirical phenomenon and examine its implications.

In recent primate work, Blanchard et al. [50] varied the post-reward delay in an ITC and found it to be systematically underestimated roughly by a factor of four, regardless of its total length (which varied across blocks from 0 to 10 seconds). What does a 4X reduction in clock speed imply about precision, and by extension, the central tendency? Should the presence of other post-reward delays in the same task significantly affect the animal’s estimates of these delays (significant central tendency)? It is not known how exactly clock speed translates to precision, but one reasonable assumption is that the clock speed and standard deviation (inverse square root of precision) scale linearly. For instance, suppose an animal learns that it should act 8 seconds after hearing a tone, which it encodes as 8 subjective seconds (e.g., 8 ticks of the internal clock), and due to timing noise, stores the interval with a granularity of 1 subjective second. This means the animal will typically respond within 7.5 and 8.5 seconds. Now imagine the internal clock were running four times slower. In that case, the animal would encode the duration as being 2 subjective seconds long (2 ticks), with a typical response occurring between 1.5 and 2.5 subjective seconds, or 6 to 10 objective (actual) seconds. Thus the standard deviation of the responses stretches by four. This in turn means that the precision will be 16 times smaller. With a large decrease in precision, our framework predicts a profound central tendency, to the point that the two posterior distributions almost overlap. Thus the animal may not discern a difference between the post-reward delays following each option. (For instance, under standard assumptions, the overlap for a 3-second and 6-second interval increases from 3% to 68%; see end of this section for a derivation.) On the other hand, if, as in the Cardinal et al. [16] result in the main text, the salience of the post-reward delay is increased, the central tendency should become less profound, and learning the contingencies should be possible. Indeed, Blanchard et al. [50] associated each option with a different post-reward delay, and found that the animals did not learn the contingencies. Furthermore, when the authors increased the salience of the post-reward delays with a small reward at the end of each one, they found that the animals’ ability to learn the option-delay contingencies was salvaged, as predicted. Note here that the posited relationship between clock speed and precision is distinct from Weber’s law, which asserts that longer intervals are more noisily encoded than shorter ones, without assuming any modifications of the clock speed [84, 89, 90].

What does the underestimation of post-reward delays imply about behavior in ITCs where the total trial length is held fixed? Here, the smaller/sooner option would have a longer post-reward delay, and the larger/later option would have a shorter post-reward delay. Notice, then, that any effect of DA on the central tendency for the post-reward delay will promote the larger/later option, which is the opposite of its effect on the pre-reward delay: With low DA, the larger/later option sees its long pre-reward delay underestimated, but its short post-reward delay overestimated, and vice versa for the smaller/sooner option. This may contribute to why DA agonists are typically found to promote the larger/later option: Both the reward and post-reward delay have a stronger central tendency than the pre-reward delay, and for both, increasing the DA level promotes the larger/later option.

Finally, it should be noted that the coupling between clock speed and precision does not qualitatively affect the DA results in the main text: A faster clock amplifies temporal estimates by the gain on the clock speed. However, this amplification would only occur if the animals were trained (i.e., learned the durations) under the faster clock. Instead, the authors administered DA agonists only after the training phase. On the other hand, the coupling does affect comparisons across the cue and no-cue conditions, and across the ascending and descending conditions. This is because, in our framework, the animal is trained on the ITCs with different temporal precisions and thus different clock speeds. Thus the trial durations of the cue and ascending conditions would be perceived to be longer than those of the no-cue and descending conditions, as they would be under the control of faster clocks. This means that, at baseline, animals in the cue and ascending conditions should favor the smaller/sooner option, as the waiting time for the larger/later option would be overestimated compared to the no-cue and descending conditions (Fig. 3I). This was indeed empirically observed, as mentioned in the main text (Fig. 3G,H).

#### Effect of encoding precision on learning post-reward delays

Under standard conditions, a 4X increase in the encoding standard deviation can result in a profound increase in the overlap between the posterior distributions for a short interval *μ_s_* and a long interval *μ_l_*. It will be convenient to consider the posterior standard deviations 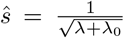. The overlap can be computed by first identifying the time *t_c_* ∈ [*μ_s_, μ_l_*] at which the posterior probabilities are equal (intersection of the two distributions):

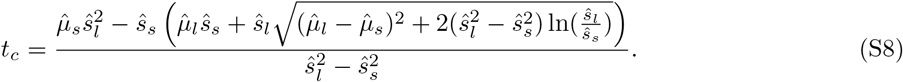

Then the overlap is

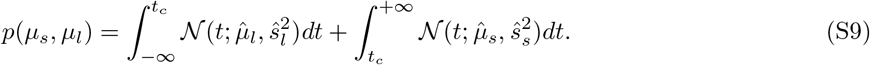

Intuitively, this represents the sum of the area under both curves to the left and to the right of *t_c_*, respectively.

For simplicity, we consider the case of a single exposure to each signal. Thus the precisions are subject to Weber’s law, which states that the standard deviations (inverse square root of precision) increase linearly with the stimulus means [84, 89, 90]. We take the Weber fraction, or the ratio of the standard deviation to the stimulus mean, to be 0.15, a typical Weber fraction for rodents in interval timing [140]. Plugging in, it follows that the overlap between the posterior distributions for a 3-second interval and a 6-second interval is 0.03 (area under the curves; maximum is 1) but increases to 0.68 when the Weber fraction is 0.15 × 4 = 0.6.

### 5 The influence of block order on temporal estimates

We have not, in this work, sought a best-fit model of the empirical data given the number of degrees of freedom available to us, including the uncertainties and their dependence on various factors (Weber’s law, repeated exposures, decay rates due to memory, capacity limitations, etc.), the shapes of the uncertainties (chosen for simplicity to be normal), the choice rule and its parameters, and so on. However, one interesting aspect of the data to model is the observed order-dependent splaying in Tanno et al. [33], wherein DA agonists cause a profound divergence of behavior for smaller delays in the descending condition. We examine this prediction here more closely.

Reinforcement learning models of interval timing tasks assume animals iteratively update their temporal estimates according to a prediction error, or the difference between the experienced and expected temporal duration:

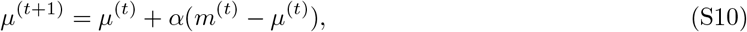

where *μ*^(*t*)^ is the animal’s estimate after the *t*-th exposure, *m*^(*t*)^ is the *t*-th (and most recently experienced) signal, and *α* ∈ [0, 1] is the learning rate. We assume that, when a new block begins, *μ*^(*t*)^ is carried over from the previous trial. Thus, with noisy learning, animals will be biased toward previously experienced intervals, across blocks.

More concretely, by sequentially expanding the second term on the right-hand side, we can rewrite Eq. (S10) as:

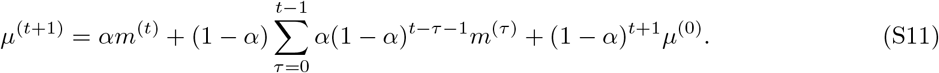

We can disregard the final term, as before experiencing the task, we assume the animal does not expect reward, i.e., *μ*^(0)^ = 0 (additionally, when *t* is large, the weight (1 − *α*)^*t*+1^ is exceedingly small). Examining the summand in the second term on the right-hand side, we see that the influence of recent history can be written as a weighted average of recent exposures, with more recent exposures being weighted more heavily than exposures that are further in the past.

Though Eq. (S10) can be rewritten to resemble Eq. (2) (i.e., *μ*^(*t*+1)^ = *αm*^(*t*)^ + (1 − *α*)*μ*^(*t*)^), it should be noted that Eq. (S10) describes *encoding* whereas Eq. (2), discussed at length in the main text, describes *decoding*. This is important because the first describes what information is stored in the brain, whereas the second describes how the animal uses this stored information to act. Acute manipulations of the estimated precision (via manipulations of DA), after learning has already occurred, do not affect the encoding process, but rather change how the encoded information is influenced by its context.

We can make two predictions from the above analysis. First, when blocks are learned in a descending order, we expect the learned temporal estimates to be biased toward larger values compared to a randomized block schedule (or one in which each block is presented on a different day). In ITCs, this means animals will be more likely to select the smaller/sooner option across DA conditions. Notice, in Fig. S1B, that the delay needs to decrease more before the animal’s choice of the larger/later option reaches the same level as in Fig. S1A, across DA conditions. Second, though temporal estimates are overestimated during learning (encoding), the central tendency during decoding, as per Eq. (2), will counteract (and possibly overwhelm) this overestimation. Thus under higher DA levels, the central tendency is masked, and the overestimation bias is more prominent. This results in a ‘splaying’ of the response curves across DA conditions (Fig. S1B).

**Figure S1:**
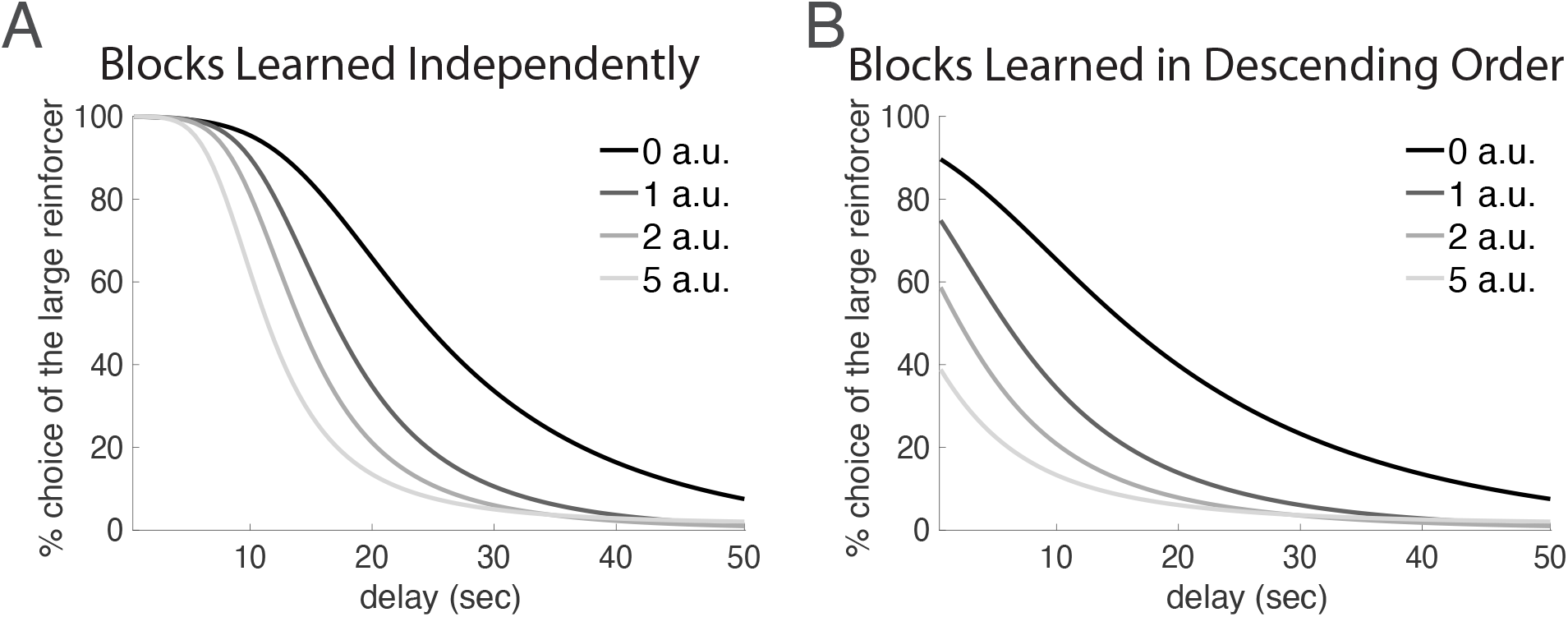
Illustration of block order influencing animal preferences. (A) We show here the typical response profile in ITCs under varying DA levels, as predicted by the Bayesian theory for the case of low temporal precision (compare with Fig. 3F). Apart from feeding low temporal precision into our model, we have not otherwise accounted for any effects of a descending block order. (B) However, when learning under uncertainty, animals will be influenced by recently experienced stimuli. Thus, when the block order of an ITC is descending, animals will tend to overestimate the temporal intervals across blocks. The temporal central tendency normally masks this overestimation, so higher DA levels will uncover it. This results in a ‘splaying’ of the response curves for smaller delays across DA levels. See Supplementary Text 3 for simulation details.

### 6 Correlating behavioral phenotypes

We have considered DA’s effects on behaviors in interval timing and ITCs. A natural extension, then, will be to examine how the behavioral phenomena covary with each other. Notably, we have predicted that higher DA should lead to more precise timing and lower apparent impulsivity in ITCs. Therefore, we predict that animals that are more precise timers should also appear less impulsive. Indeed, Marshall et al. [98] examined rats’ impulsivity and timing abilities. To assess their impulsivity, the authors trained the rats on a standard ITC. To assess their timing precision, the authors trained the rats on a bisection task [141]: Here, the rats were trained to respond with, for example, a left lever press when presented with a short (4-second) interval, and with a right lever press when presented with a long (12-second) interval. They were then tested on intermediate-duration intervals, for which they could still only respond with either a left lever press (the short-duration response) or a right lever press (the long-duration response). The stochasticity of responses was taken to reflect timing noise. The authors found that the more precise timers also tended to be less impulsive (Fig. S2A), as predicted by our framework (Fig. S2B). McClure et al. [99] also examined the correlation between timing precision and impulsivity, but using a peak-interval task, in which animals are trained to *reproduce* experienced durations, rather than a bisection task, in which animals are trained to *estimate* them, and reported similar findings.

**Figure S2:**
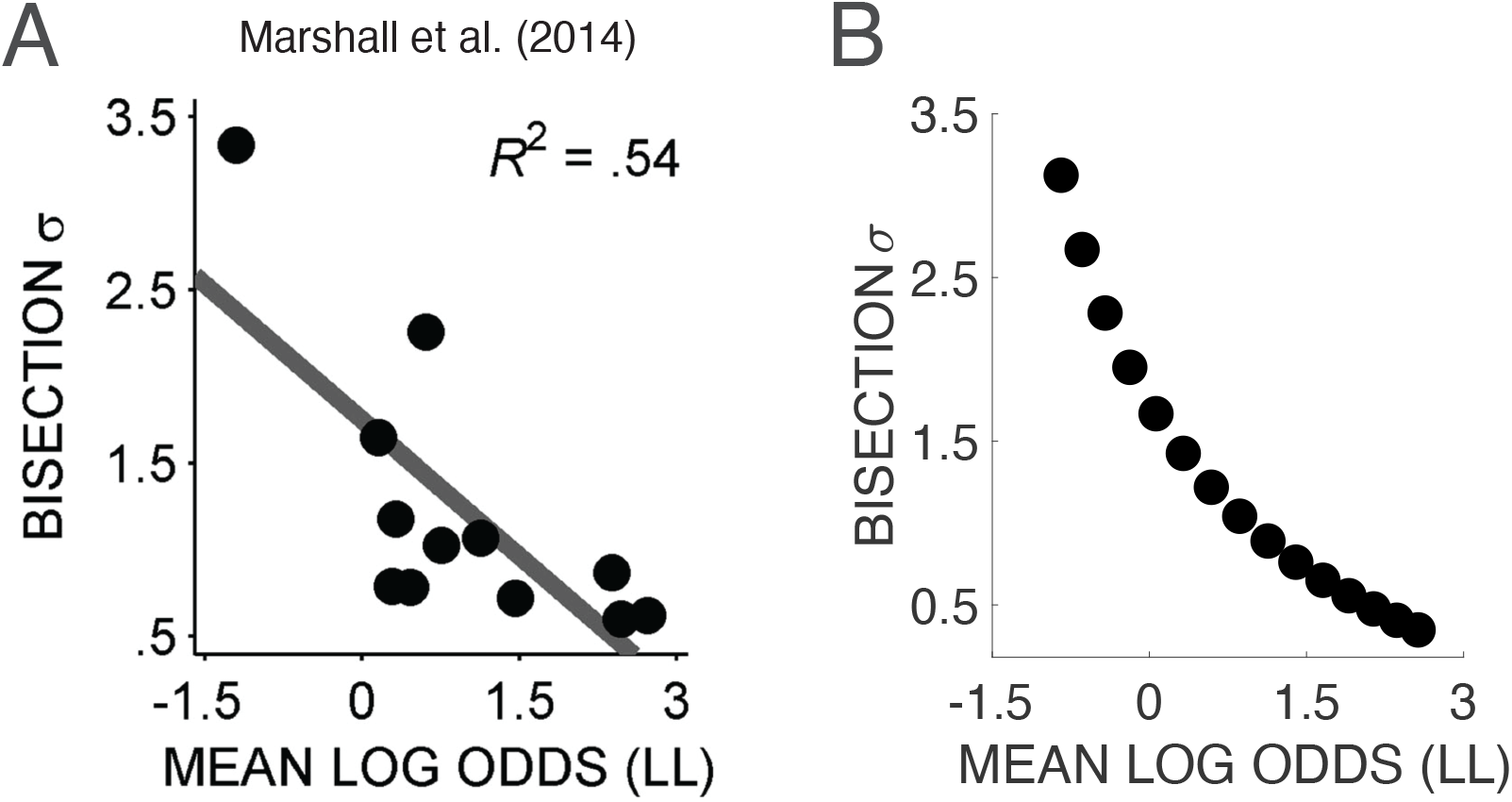
More precise timers are more likely to select the larger/later option. (A) Marshall et al. [98] measured rats’ temporal precision and ‘impulsivity’ using a bisection task and ITC, respectively. The authors found that lower noise in the bisection task correlated with a tendency to select the larger/later option. Mean log odds are defined as the mean logarithm of the ratio of the probability of selecting the larger/later option to the probability of selecting the smaller/sooner option. LL: larger/later option, *σ*: parameter fit to computational model representing stochasticity of choices. (B) Our model recapitulates this effect: Animals with higher DA levels are predicted to display more precise timing and a tendency to select the larger/later option. See Supplementary Text 3 for simulation details.

Interestingly, Marshall et al. [98] also examined the relationship between impulsivity and reward magnitude sensitivity, which they studied using a two-armed bandit task where the larger reward was varied across blocks. The authors did not find a relationship between the two, although, as they note, this may be due to an inadequate metric for quantifying reward sensitivity (ratio of large-reward lever press rate to the sum of large- and small-reward lever press rates).

### 7 Dopamine and the rat gambling task

Having considered delay discounting and probability discounting separately, let us now examine the rat gambling task (rGT), which involves a manipulation of both the delays and the reward probabilities.

The rGT, adapted from the Iowa gambling task [142], is a four-armed bandit task of fixed block duration, where each arm yields a reward of different magnitude, delivered with a different probability, and where omissions are accompanied by ‘punishing’ time-outs of different durations (Table S1). The higher the reward magnitude, the lower the probability of reward and the longer the time-out during omissions [105].

**Table S1:**
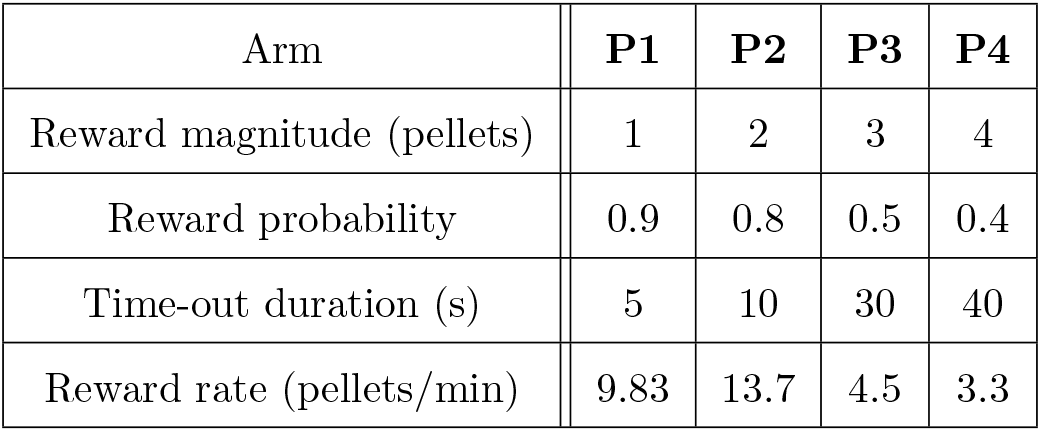
Properties of each arm in the rGT. The advantageous arms *P*1 and *P*2 yield relatively small rewards with high probability and include a short time-out duration during reward omissions. The disadvantageous arms *P*3 and *P*4 yield relatively larger rewards with lower probability and include a longer time-out duration during reward omissions. Defining *p* as the probability of obtaining reward, the probability of an omission and time-out is (1 − *p*). Let *A* be the inter-trial interval, which in Zeeb et al. [105] is 5 seconds. This means that the duration of a trial in which a time-out *t* occurs is *A* + *t*. The reward rate is then 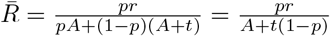.

The arms *P*1 and *P*2 in Table S1 are referred to as the ‘advantageous arms’ because they yield high reward rates [143]. They are characterized by relatively small reward magnitudes but high reward probabilities and short time-out durations during reward omissions. On the other hand, *P*3 and *P*4 are the ‘disadvantageous arms.’ They yield low reward rates and are characterized by relatively large reward magnitudes, low reward probabilities, and short time-out durations. The reward rate is highest for *P*2 and lowest for *P*4; thus, with perfect learning, rats should preferentially select *P*2 and be most averse to selecting *P*4.

To study the dopaminergic modulation of risk-seeking behavior, Zeeb et al. [105] began by training rats on the rGT. The authors found that the rats were indeed far more likely to select *P*2 than any other arm. However, the tendency to select *P*4—objectively yielding the worst reward rate—was higher than that of *P*3 (Fig. S3A), a peculiar but well-replicated finding in rGTs [143]. After the animals achieved stable behavior, the authors administered the DA agonist amphetamine and tested them again on the task. While the rats still displayed the same qualitative preferences, they became more likely to select *P*1 and less likely to select *P*2 compared to baseline (Fig. S3A), which was taken to indicate that the DA agonist induced a decline in performance (see also [144] who observed this effect in mice).

**Figure S3:**
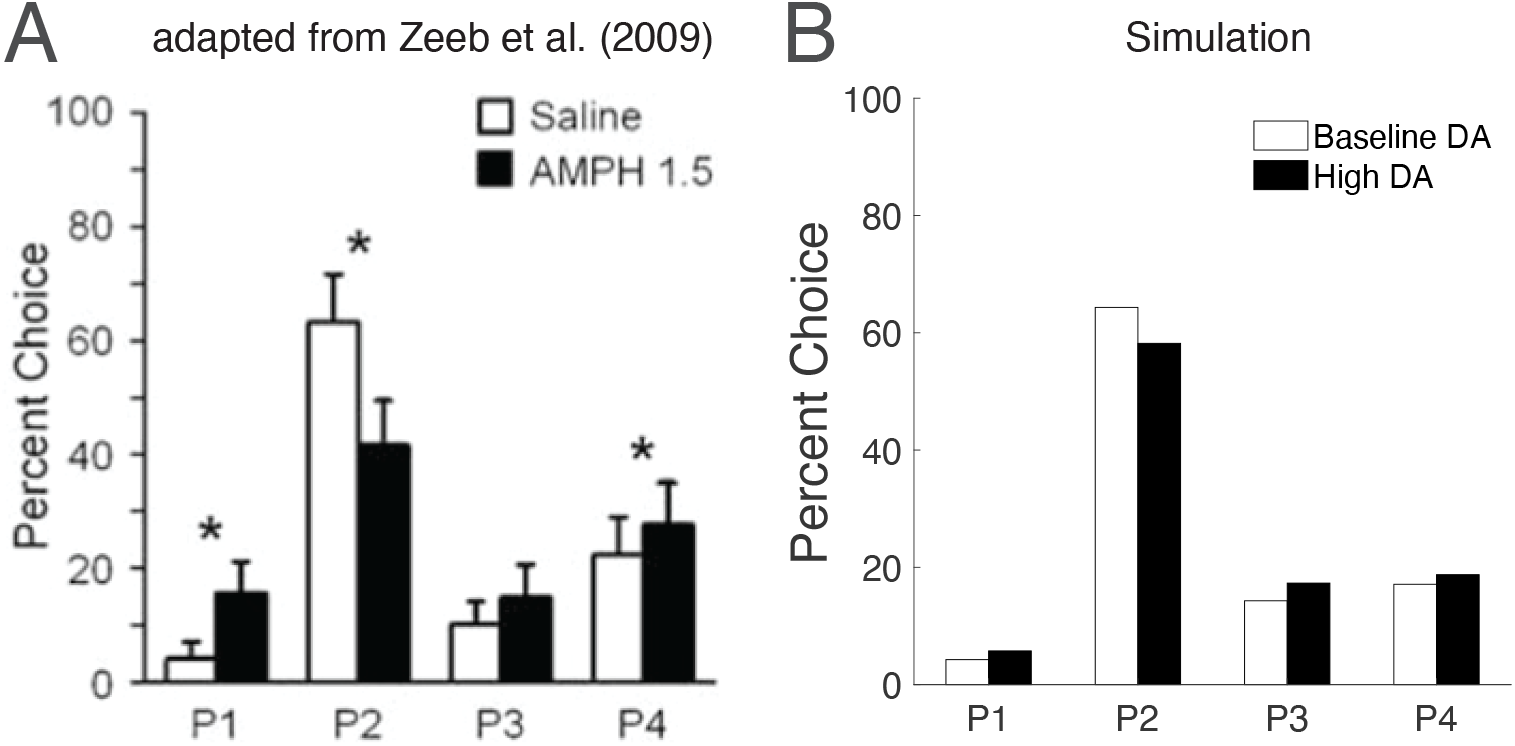
The Bayesian theory and the rGT. (A) Zeeb et al. [105] trained rats on an rGT that included four arms, each yielding a different reward with a different reward probability and a different time-out duration during omissions (Table S1). The authors found that animals prefer arm *P*4 to *P*3 at baseline, even though *P*4 yields fewer rewards over the course of a fixed 30-minute block. Administration of the DA agonist amphetamine promotes *P*1 and suppresses *P*2. (B) The Bayesian theory captures these effects. We have assumed a hierarchical Bayesian model where the advantageous arms *P*1 and *P*2 form one context, the disadvantageous arms *P*3 and *P*4 form another, and all four arms form the hyper-context. With a strong temporal central tendency, the longest duration, corresponding to *P*4, is underestimated and the second-longest duration, corresponding to *P*3, is overestimated. This benefits *P*4 over *P*3. With higher DA levels, the temporal central tendency is silenced, so the estimated duration corresponding to *P*1 decreases and that corresponding to *P*2 increases. This benefits *P*1 over *P*2. See Supplementary Text 3 for simulation details.

We will show that both of these findings can be accommodated by the Bayesian framework. Importantly, we will begin by seeking to limit the examined parameter space, as the Bayesian framework becomes very flexible with four signals in each domain. To illustrate this flexibility, recall that, in ITCs and PDs, it was enough to examine the relative strength of each central tendency, and the encoding precisions of each arm were otherwise irrelevant. This is because increasing the encoding precision for either the small or large reward yields the same effect—an amplification of the difference in posterior means (black segments in Fig. 1A,B). On the other hand, with four rewards, increasing the encoding precision for one may increase the estimated difference between some pairs but decrease it between others. A second difficulty is that the context itself is ill-defined. Generally, contexts defined over multiple signals are better captured by hierarchical Bayesian frameworks in which different levels of the hierarchy have different priors. This is the case for the temporal central tendency, where intervals within ‘mini-contexts’ are attracted toward their own prior means, which in turn are attracted to a hyper-prior mean over a larger context [75]. For the rGT, the properties of *P*1 and *P*2 are far more similar to each other than to *P*3 or *P*4 (indeed, the four arms are divided into ‘advantageous’ and ‘disadvantageous’ arms), and therefore the context for *P*1 may consist largely of the advantageous pair. To limit this flexibility, we will make two assumptions: first, that the temporal central tendency dominates the reward central tendency. This is in line with previous work that has found post-reward delays and inter-trial intervals to be encoded with profoundly low temporal precision [50]. As the omission durations drive the trial length in this task, we find this to be a reasonable assumption. Second, we will assume a hierarchical Bayesian framework in which the advantageous arms form one context, the disadvantageous arms form another, and the four arms together form the hyper-context. This will allow for an intuitive explanation of the two observed phenomena.

Let us begin with the *P*4 finding. When the temporal central tendency is strong, the delay of *P*4 will be underestimated, so its estimated reward rate will increase. Within the context of disadvantageous arms, the delay of *P*3 will instead be overestimated. Thus the temporal central tendency will benefit *P*4 compared to *P*3. Examining now the context of advantageous arms, the delay of *P*1 is normally overestimated and that of *P*2 is underestimated. DA agonist administration silences the temporal central tendency, which benefits *P*1 compared to *P*2. Thus if the temporal central tendency is sufficiently strong at baseline, the animal will prefer *P*4 to *P*3, and will increase its preference for *P*1 but decrease it for *P*2 with the DA agonist (Fig. S3B), as observed.

We have shown that the finding that *P*4 is selected more often than *P*3, even though it yields the lowest reward rate, can be accommodated by the Bayesian framework. The Bayesian theory of DA furthermore captures the finding that DA agonists promote selection of *P*1 and inhibit selection of *P*2. Note, however, that these results depend on the chosen parameter space, owing to the flexibility of the Bayesian theory in this task (i.e., the number of its degrees of freedom). The benefit of the theory is that it can account for empirical phenomena that otherwise require additional assumptions about animal preferences and the roles of DA.

